# Polar tube firing dynamics and ultrastructure from the shrimp microsporidian *Ecytonucleospora hepatopenaei* (EHP)

**DOI:** 10.1101/2025.07.23.666298

**Authors:** Nutthapon Sangklai, Thanamon Udomphonphaibun, Krittapron Keawpanya, Paris R. Watson, Nicolas Coudray, Maia A. Koliopoulos, Orawan Thepmanee, Arun K. Dhar, Damian C. Ekiert, Gira Bhabha, Anchalee Tassanakajon, Pattana Jaroenlak

## Abstract

*Ecytonucleospora (Enterocytozoon) hepatopenaei* or EHP is an obligate intracellular parasite that belongs to phylum Microsporidia. EHP infection in shrimp results in growth retardation and size variation, leading to severe economic loss to shrimp aquaculture worldwide. Similar to other microsporidian species, EHP utilizes a harpoon-like invasion apparatus called the polar tube in order to infect host cells. The polar tube typically coils inside the spore and rapidly transits to a long, extended tube after being activated with proper stimuli. However, the mechanism and physical conditions affecting the polar tube firing in EHP are poorly understood. Here, we screened several germination buffers and found that a buffer containing potassium hydrogen phthalate (KHP) at pH 3.0 yielded the highest germination rate. The optimal temperatures for EHP germination range between 25-28°C, similar to the shrimp’s body temperature. The spores require at least 30 seconds to be activated, suggesting that the stimuli could rapidly move across the spore wall. We utilized high-speed live-cell imaging to study the dynamics of the EHP polar tube firing and compared the dynamics when the spores were treated with KHP and a previously reported stimulus, phloxine B. The polar tube firing dynamics between these two conditions are different. The total firing time was ∼100 milliseconds with the maximum velocity of ∼300 μm⋅s^-1^ in KHP condition. Further investigation on the EHP polar tube ultrastructure using cryo-electron microscopy revealed that the EHP polar tube was composed of a membrane layer with an additional repetitive protein-array on the outermost surface. The distance between each repetitive unit ranges between 52-66 Å. These repetitive units could possibly be polar tube proteins (PTPs). Altogether, this study provides insights into EHP polar tube firing dynamics and its architecture which are important foundations for understanding the biophysical factors governing EHP pathogenesis and in developing EHP control strategies.

**Author summary:** The microsporidian *Ecytonucleospora hepatopenaei* (EHP) poses a significant threat to global shrimp aquaculture, causing substantial economic losses. Many studies on EHP have been focused on development of detection methods, and farm management. Little is known about EHP pathogenesis and how it infects shrimp cells. Here, we identified laboratory conditions to activate the parasite’s infection process, which occurs on a millisecond timescale. We utilized a high-speed live-cell imaging to further characterize the rapid EHP infection process. The architecture and ultrastructure of EHP invasion organelle were also investigated using cryo-electron microscopy. Our findings provide a better understanding of the EHP infection mechanisms that are crucial for developing effective control strategies.

## Introduction

The microsporidian *Ecytonucleospora (Enterocytozoon) hepatopenaei* or EHP is an economically important parasite in shrimp aquaculture. EHP is the causative agent of hepatopancreatic microsporidiosis (HPM) [1]. The primary symptom of HPM is growth retardation, which results in size variability among shrimp [2]. Even though severe EHP infections do not lead to mortality, they significantly decrease shrimp farm productivity, resulting in major economic loss [3,4]. Infection with EHP also leads to susceptibility to secondary infections by bacteria and viruses [5,6]. The primary infection site of EHP is tubule epithelial cells of the hepatopancreas [2]. Since hepatopancreas is an important organ for production of several digestive enzymes, digestion of food and absorption of nutrients, EHP infected shrimp experience abnormal, retarded growth [7,8].

Typically, EHP and other microsporidian species develop environmentally resistant spores as a part of their life cycles. The spores are equipped with a specialized infection organelle called the polar tube [9–11]. Under suitable conditions or when the spores encounter host cells, the spores rapidly fire the polar tube. This process is called spore germination or polar tube firing [12,13]. The polar tube potentially penetrates host cell membranes and serves as a conduit to transfer infectious contents into the host [14]. Laboratory conditions to trigger spore germination greatly among microsporidian species, for example *Encephalitozoon* spp. can be triggered by hydrogen peroxide (H_2_O_2_) [14], while dehydration and pH shift are common triggers for *Nosema* spp. [14]. For EHP, a red, water-soluble dye, called phloxine B has been reported to initiate the polar tube firing [15]. Many studies have shown important factors that are essential for spore germination such as shift of the pH [16,17], monovalent cations [18], and salt concentrations [18]. It is important to note that common factors that share among microsporidian species are not known since there is no systematic study testing these conditions in different microsporidian species. Hence, the mechanisms underpinning the polar tube firing are largely unknown and not yet generalizable.

Among different factors that trigger for the polar tube firing, pH appears to be the most common factor. An optimal pH is required to yield the highest percentage of spore germination [16,19]. In many microsporidian species, a shift from a neutral pH to a basic pH is sufficient to trigger spore germination. For example, shifting of 1-2 pH units activates germination of *Glugea hertwigi* [20]. In contrast, pH shift from neutral to acidic condition is required for polar tube firing in *Nosema pulicis* [21] and *Pleistophora anguillarum* [22]. In addition, it has been reported that polar tube firing in *Vavraia culicis* is pH independent since the spores can be activated in both acidic and basic conditions [23]. Despite these findings, the role of the pH alone is unclear. It is generally studied together with other stimuli such as ions.

Alkali metal ions such as Na^+^ and K^+^ are another important factor for spore germination. Several studies tested the capacity of metal ions to trigger spore germination [18,24,25]. In *Anncaliia algerae* (formerly of the genus *Nosema*), the efficiency of alkali metal ions follows K^+^ > Na^+^ > Rb^+^ > Cs^+^ > Li^+^ in order of decreasing efficiency [25]. Substitution of Na^+^ and K^+^ with a membrane impermeable cation, choline chloride, completely inhibits spore germination [18]. The germination was restored when Na^+^ and K^+^ were added back into the buffer. These results suggest that cations must enter into the spore to activate spore germination. Effects of Na^+^ and K^+^, and other alkali cations on spore germination vary among microsporidian species.

The process of polar tube firing happens extremely fast on a millisecond timescale [12]. Due to its fast nature, it is challenging to study the germination process. Recent works have utilized high-speed live-cell imaging to better understand the kinetics of polar tube firing in four microsporidian species that infects mosquitoes and humans, including *A. algerae*, *Encephalitozoon hellem*, *Encephalitozoon intestinalis* [26], and *Vairimorpha necatrix* [27]. The polar tube firing can be divided into 3 phases, including 1) polar tube elongation, 2) polar tube static phase, and 3) emergence of the infectious material or sporoplasm [26]. The fired polar tube experiences high acceleration to reach a speed of ∼230 μm⋅s^-1^ in *A. algerae* and 330 μm⋅s^-1^ in *E. hellem*, possibly generating enough forces to pierce host cell membranes [26,28]. From this kinetics study, variation between species is clearly observed, even between closely related species *E. hellem* and *E. intestinalis* [26].

Little is known about the physical conditions affecting the EHP spore germination as well as the EHP polar tube firing kinetics. Here, we investigated the effects of pH, monovalent cations, temperature, and activation time on the EHP spore germination. High-speed live-cell imaging was used to study the dynamics of EHP polar tube firing and to observe the infectious cargo transport during the spore germination. We further visualized the extruded EHP polar tube using cryoEM. This study provides insights into EHP infection mechanisms. Germination conditions identified in this work could be further tested to develop anti-EHP agents, which will minimize the impact of EHP in farmed shrimp.

## Results

### Physical conditions affecting EHP spore germination

Since phloxine B is the only chemical that has been reported to trigger EHP spore germination [15], here we screened panels of germination buffers, ranging from pH 2.0-14.0 (See Methods, S1 Fig). The results showed that EHP spores could germinate in two preferred pH windows (Fig. 1A). Interestingly, EHP spores preferentially germinate under acidic pH. The germination buffer containing 100 mM potassium hydrogen phthalate (KHP) at pH 3.0 showed the highest germination rate at 85.7 ± 2.5%, compared to ∼18% using the ammonia-ammonium chloride buffer at pH 9.0 (Fig. 1A). Considering that lower pH yields higher EHP germination rate, we further tested another germination buffer at pH 3.0. The results showed that the citrate buffer, pH 3.0 failed to trigger the EHP spore germination (S1 Table). We also tested phthalate buffers at other pH, including pH 2.0 and 4.0. Both conditions failed to trigger spore germination. Hence, our results suggest that both phthalate and pH 3.0 contribute to the EHP spore germination, *in vitro*.

**Fig. 1.**
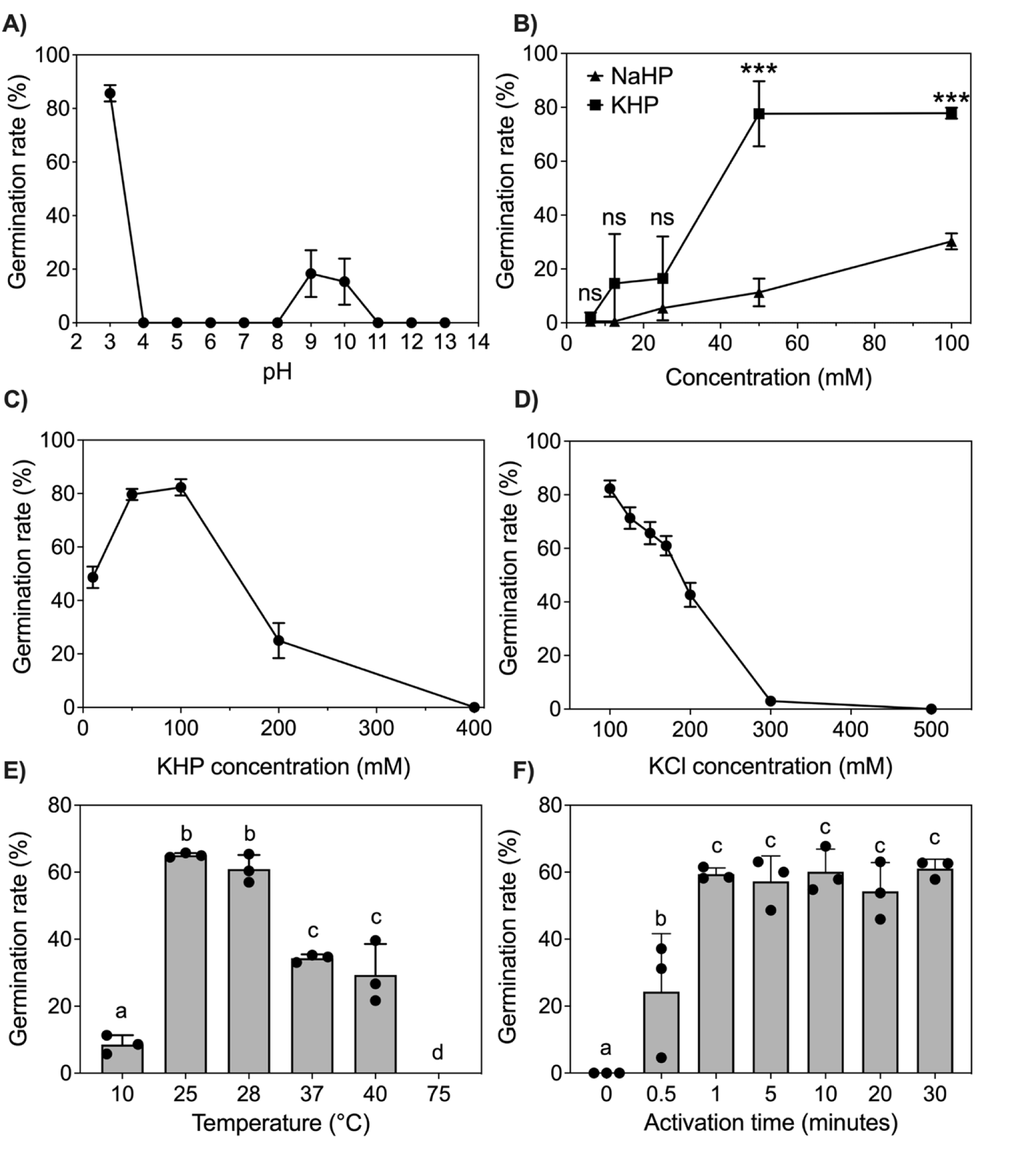
Effect of physical conditions on the EHP spore germination. Quantification of the EHP spore germination rate when the spores were germinated in different (A) pH and germination buffer, (B) monovalent cations, (C) KHP buffer concentrations, (D) KCl concentrations, (E) temperatures, and (F) activation time. Each experiment was performed in three biological replicates (n=100 for each replicate). Error bars represent the standard deviation of three biological replicates. A one-way ANOVA was used to compare the significance of monovalent cations, *** indicates P<0.001. Student t-test was used to determine the significance of temperature and activation time. In (E) and (F), a– d indicates significant differences among groups (P<0.05).

It has been widely accepted that monovalent cations impact microsporidian spore germination and each microsporidian species prefers different monovalent cations [12]. Here, we tested the effect of different monovalent cations on EHP spore germination using potassium hydrogen phthalate (KHP) and sodium hydrogen phthalate (NaHP) buffers, while maintaining the buffer concentration at 100 mM and the pH of both solutions at 3.0. The results showed that the EHP germination rate was ∼2.7 times higher in KHP than NaHP (77.8% ± 1.7% vs 28.2% ± 1.5%) (Fig. 1B), suggesting that EHP spores prefer to germinate under K^+^ more than Na^+^ condition. It is possible that K^+^ has a smaller hydrated ion radius than Na^+^ ion [29], hence it better penetrates through the EHP spore wall and triggers spore germination. These results led us to hypothesize that if we increase the concentrations of K^+^ or phthalate, they would enhance the EHP spore germination. To test this hypothesis, we first varied KHP buffer concentrations while maintaining the pH of the buffer at 3.0. The results showed that the optimal germination buffer concentrations ranged between 50 mM and 100 mM (Fig. 1C). Additional increases in buffer concentrations reduced the EHP germination rate and completely inhibited the EHP germination at 400 mM concentration (Fig. 1C). The inhibitory effect was consistent when we additionally applied KCl to the 100 mM KHP buffer (Fig. 1D). It is possible that concentrations higher than the optimal concentrations would be considered a hypertonic solution. Hence, the EHP spores would lose water that is retained inside the spore and is required for a successful spore germination [30].

Normally, the body temperature of the shrimp is approximately 25-30°C [31]. To test whether these are optimal temperatures for the EHP spore germination, *in vitro*, we germinated the EHP spores at various temperatures, including 10, 25, 28, 37, 40, and 75°C using a 100 mM KHP, pH 3.0 buffer. As expected, 25 and 28°C yielded the highest germination rates (Fig. 1E), resembling the actual condition inside the shrimp host. In addition, incubating the EHP spores at 75°C completely inhibited spore germination (Fig. 1E), similar to a previously published study [32].

Next, we investigated how long it takes for the EHP spores to germinate after being incubated with the germination buffer. Our results showed that the spores begin germinating rapidly, less than 30 sec after being in the germination buffer, and most germination events were completed within 1 min (Fig. 1F). This implies that the stimulants would quickly move across a ∼50-nanometer thick spore wall layer on a fast timescale (in less than 30 sec) [33].

After the polar tube is being fired from the microsporidian spores, sporoplasm or infectious cargo is transported into host cells and propagates into new parasites [26,34]. To examine whether our *in vitro* germination conditions yield active sporoplasm, we observed the presence of the sporoplasm at the end of the polar tubes and used it as an indicator for active sporoplasm (Fig. 2A and 2B). The EHP spores were germinated with 2% phloxine B or 100 mM KHP, pH 3.0. The results showed that germination with KHP yielded a significantly higher percentage of sporoplasm occurrence than 2% phloxine B (Fig. 2C and 2D). This implies that KHP triggers a more complete polar tube germination and this condition is suitable for characterization of the polar tube firing kinetics.

**Fig. 2.**
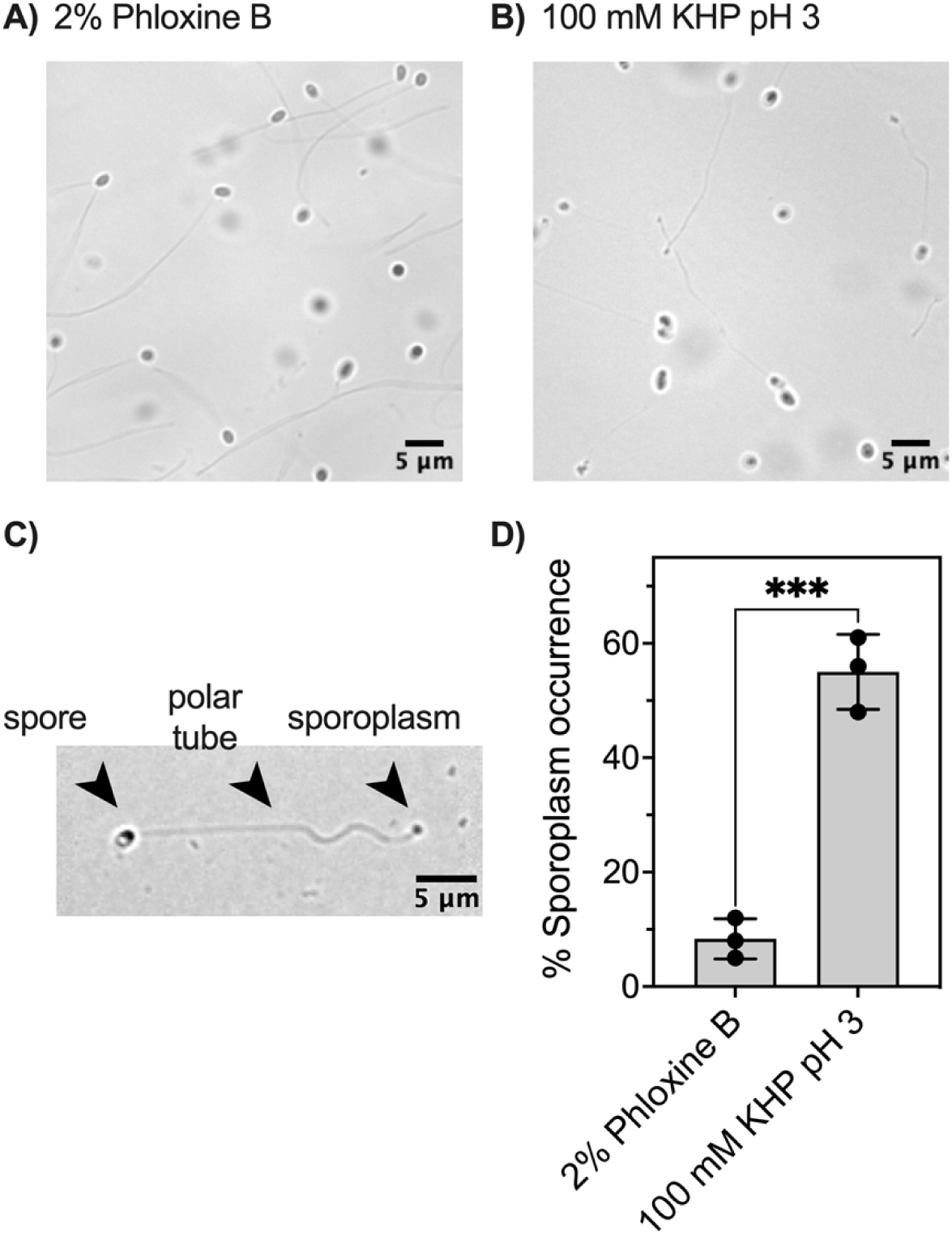
Quantification of the EHP sporoplasm after being induced with different germination buffers. The spores were induced with (A) 2% phloxine B and (B) 100 mM KHP, pH 3.0. (C) A representative phase contrast micrograph showing sporoplasm at the distal end of the polar tube. (D) Percentage of the sporoplasm occurrence. Statistical significance was determined by a student t-test (*** P<0.001).

Taken together, the optimal, *in vitro* conditions for the EHP spore germination follows: the spores are germinated with a 100 mM KHP at pH 3.0 for 1 min at either 25°C or 28°C. Our results provide better understanding on the EHP *in vitro* germination conditions that possibly mimics the actual conditions within shrimp hosts.

### Kinetics of the EHP spore germination

Polar tube firing process in microsporidia is one the most fascinating and extremely rapid processes in biology [28] This process takes place on a millisecond timescale (in less than 1 sec), making it challenging to image in real time [26]. Previously, a high-speed live-cell imaging has been used to study the polar tube firing kinetics in four microsporidian species, including two species of human-infecting microsporidian and two species that infect insects [26,27]. However, the polar tube firing kinetics in microsporidian species infecting aquatic animals remains largely unexplored. Here, we compared the kinetics of the EHP polar tube firing when triggered with 100 mM KHP pH 3.0 and 2% phloxine B. With a frame rate of ∼170 fps, we captured the entire EHP polar tube firing process from 20 different spores per condition (Fig. 3A and 3B, S1 and S3 Videos). In the KHP condition, two distinct phases were observed, namely the polar tube elongation phase (phase 1), and the emergence of sporoplasm (phase 2) (Fig. 3A, S1 Video). Interestingly, we could not observe the polar tube static phase, which was previously reported in *A. algerae*, *E. intestinalis,* and *E. hellem* [26]. Next, we quantified the kinetics of the EHP polar tube firing when triggered with KHP, including polar tube length, time used to extend to ∼90% of the total lengths (T_EXT90_), and maximum velocity (V_max_) (Fig. 3C-3F). The results showed that the EHP germination process takes place in 116.8 ± 26.8 ms (Fig. 3E), extruding the average polar tube length of 25.8 ± 3.4 μm (Fig. 3C and 3D) with the V_max_ of 311.5 ± 111.3 μm⋅s^-1^ (Fig. 3F). It is important to note that these polar firing kinetics are indistinguishable when the spores were triggered with KHP, at either 25°C or 37°C (Fig. 3C-3F, S2 Video).

**Fig. 3.**
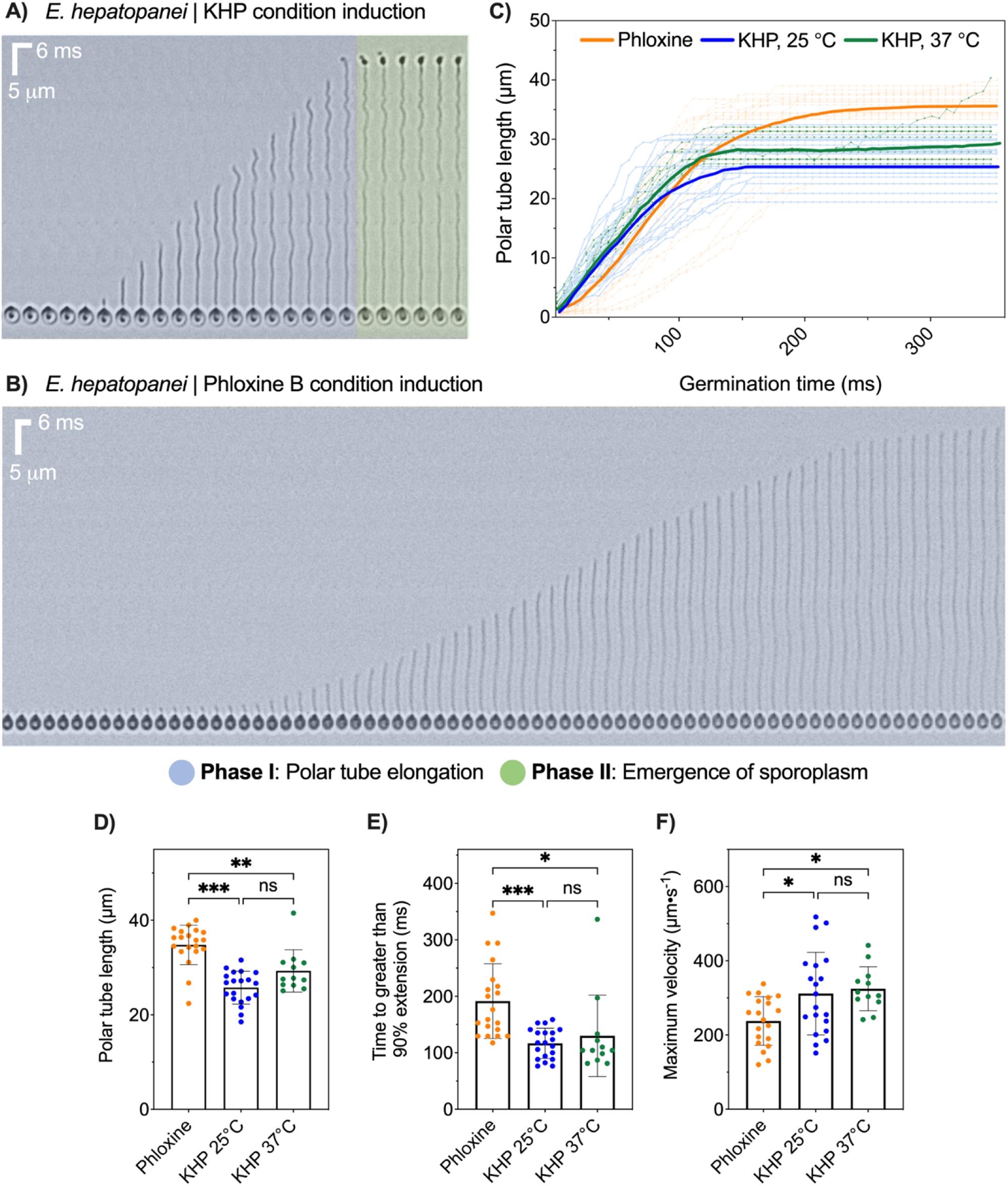
Polar tube firing kinetics of the EHP spores. (A) A representative kymograph showing the spore germination after being induced with (A) 100 mM KHP pH 3.0, and (B) 2% phloxine B. (C) Quantification of the polar tube length as a function of time from three different germination conditions, including 2% phloxine B, 100 mM KHP at 25°C, and 100 mM KHP at 37°C. (D) Quantification of the polar tube length and (E) time taken for the polar tube to extend to ≥90% of its maximum length (TEXT_90_). (F) Average maximum velocity of the polar tube extension process. Note that n = 20 for phloxine and KHP at 25 °C, while n = 12 for KHP at 37 °C in all graphs. All the data were analyzed by one-way ANOVA. The error bars in this figure represent standard deviations. * P<0.05, ** P<0.01, and *** P<0.001.

In contrast to KHP, 2% phloxine yields altered polar tube firing kinetics (Fig. 3C-3F). First, only the polar tube elongation phase (phase 1) was observed. In all 20 tested spores, we could not observe the emergence of the sporoplasm at the end of the tube (Fig 3B, S2 Video). Quantification of the polar tube firing kinetics showed that the polar tube length is ∼10 μm longer than those triggered by KHP (Fig. 3C and 3D). In addition, EHP spores take ∼75 ms longer to germinate (T_EXT90_ = 191.5 ± 64.5 ms) with a lower speed (V_max_ 237.6 ± 64.0 μm⋅s^-1^) when germinating under the phloxine B condition (Fig. 3E). It is unclear why these two germination buffers result in different polar tube firing kinetics, even though the spores were prepared from the same EHP-infected hepatopancreas. It is plausible that these two buffers alter protein conformations and organizations on the polar tubes.

### Visualization of the EHP polar tube using cryo-EM

To test whether the overall appearance and ultrastructure of the EHP polar tube differs when triggering with different germination buffers, we visualized the EHP polar tube using cryo electron microscopy (Fig. 4A, 4B). Our results showed that the overall polar tube morphology is similar between the KHP and phloxine B induced spores (Fig. 4A, 4B). 2D class averaging analysis showed that the EHP polar tube comprises a membranous layer outside of which is another layer that consists of repeating densities, likely protein (Fig. 4C). This is consistent with previous observations by cryo-EM in *N. bombycis* [35]. Measuring the thickness of the membranous layer showed an average between 23-32 Å (S3 Fig and S4 Fig), consistent with biological membranes [36–38]. For the outermost layer, we hypothesized that this layer could be a protein-array and made of repetitive protein subunits. Quantification of the distance between the center of the repetitive protein unit to the next showed the distance ranging between ∼52-66 Å (Fig 4D, 4E). The distance of this repetitive protein unit found in EHP is similar to those reported in the outer filament (OF) layer of the *E. intestinalis* polar tube [39]. While the proteins that form the core structure of the polar tube have not been well defined, a few polar tube proteins (PTPs) have been hypothesized to form the structural basis of the polar tube [40–42]. It is possible that the density we observe here represents one or more of the PTPs. So far, only PTP2 has been characterized in EHP [43]. Further proteomic study needs to be performed in order to identify proteins that make up the EHP polar tube.

**Fig. 4.**
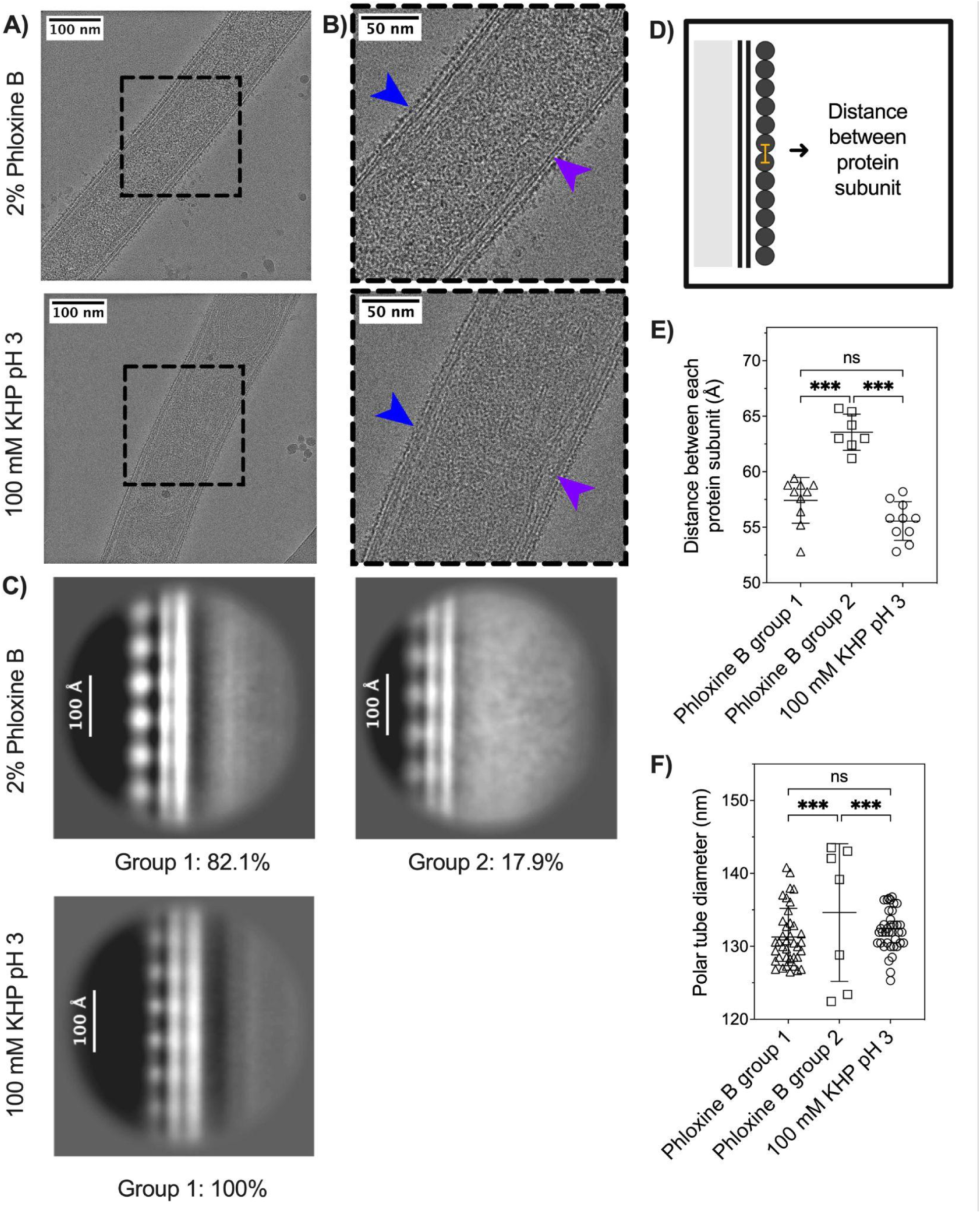
Visualization of the EHP polar tube using cryo-EM. (A) Representative Cryo-EM micrographs of the polar tube after being germinated with phloxine B (upper panel) and KHP (lower panel). The scale bars are 100 nm. (B) Zoomed-in images from (A). Blue arrowheads indicate the outermost repetitive protein layers, while purple arrowheads represent the membrane layer. (C) Representative 2D class averages of the polar tube from each germination buffer (See S2 Fig for gallery of all classes). (D) A diagram showing how the distance between the protein subunits was measured. (E) Quantification of the distance between each repetitive density on the outermost layer of the tube. (F) Quantification of the polar tube diameter. Statistical significance was determined by one-way ANOVA (*** P<0.001, ** P<0.05). Error bars represent SD, n_phloxine B group 1_ = 10, n_phloxine B group 2_ = 7, n_KHP_ = 10 in E and F. In G, n_phloxine B group 1_ = 38, n_phloxine B group 2_ = 7, nKHP = 48.

Further analyses of our 2D class averaging of the EHP polar tube revealed two subclasses when the tube was triggered by the phloxine B (Fig. 4C), which differed in the appearance of the repeating density in the outermost layer of the tube. From the original 45 micrographs, group 1 constitutes 1,676 particles (or 82.1%), while group 2 contains 376 particles (17.9%) (Fig. 4C, S2 Fig). The basis for the different populations is not clear, but we observed that particles in groups 1 and 2 typically originated from different polar tubes. This suggests that either the polar tubes are in different stages of firing, or there might be heterogeneity in the polar tube morphology. In only 10% of the micrographs, we found particles from both groups on the same polar tube. In contrast, only one class of the tube was found in the KHP condition, which contains a total of 2,174 particles (Fig. 4C and corresponds to group 1 for the phloxine B condition). Comparison of the tube ultrastructure between these subgroups showed that the distance between each repetitive protein subunit (Fig. 4E) and the polar tube diameter (Fig. 4F) are indistinguishable between phloxine B group 1 and KHP groups. However, these two parameters are slightly higher when compared to the phloxine B group 2 (Fig. 4E, 4F). It is unclear whether these differences will contribute to the differences in the polar tube firing kinetics observed in Fig. 3F since phloxine B group 2 constitutes only ∼18% of the entire phloxine B population.

### Infectious cargo transport through the polar tube

One function of the polar tube is to provide the conduit for the infectious cargo transport. The infectious cargo consists of the parasite’s nucleus and ribosomes, important for successful infection and proliferation [26,44]. Previous work attempted to elucidate the dynamics of cargo transport by tracking the parasite’s nuclei using a fluorescent dye, NucBlue in mosquito-infecting microsporidian *A. algerae* [26]. Applying a similar technique, we labeled the EHP nucleus and observed its transport through the polar tube (Fig. 5A, 5B). Using duo-detection of fluorescence and transmission light, we found that the EHP polar tube firing takes place prior to the nuclear translocation, similar to those in *A. algerae* [26] (Fig. 5B, S4 Video). The EHP nuclear transport begins ∼66 ms after the polar tube is fired (Fig 5B). This is the point where the polar tube is extruded to approximately 50%. Consistent with *A. algerae*, massive nuclear deformation is observed in EHP (Fig 5C, S5 Video). The EHP nucleus travels at the rate of ∼288 μm⋅s^-1^ (Fig. 5D). It is important to note that we were not able to track the entire nuclear translocation from the spore body to the distal end of the polar tube. From all the videos obtained, nuclear transport is either incomplete or the nucleus goes out of the imaging field. Our results suggest that the nuclear transport process seems to be conserved among different microsporidian species, or at least between EHP, *N. bombycis* and *A. algerae*.

**Fig. 5.**
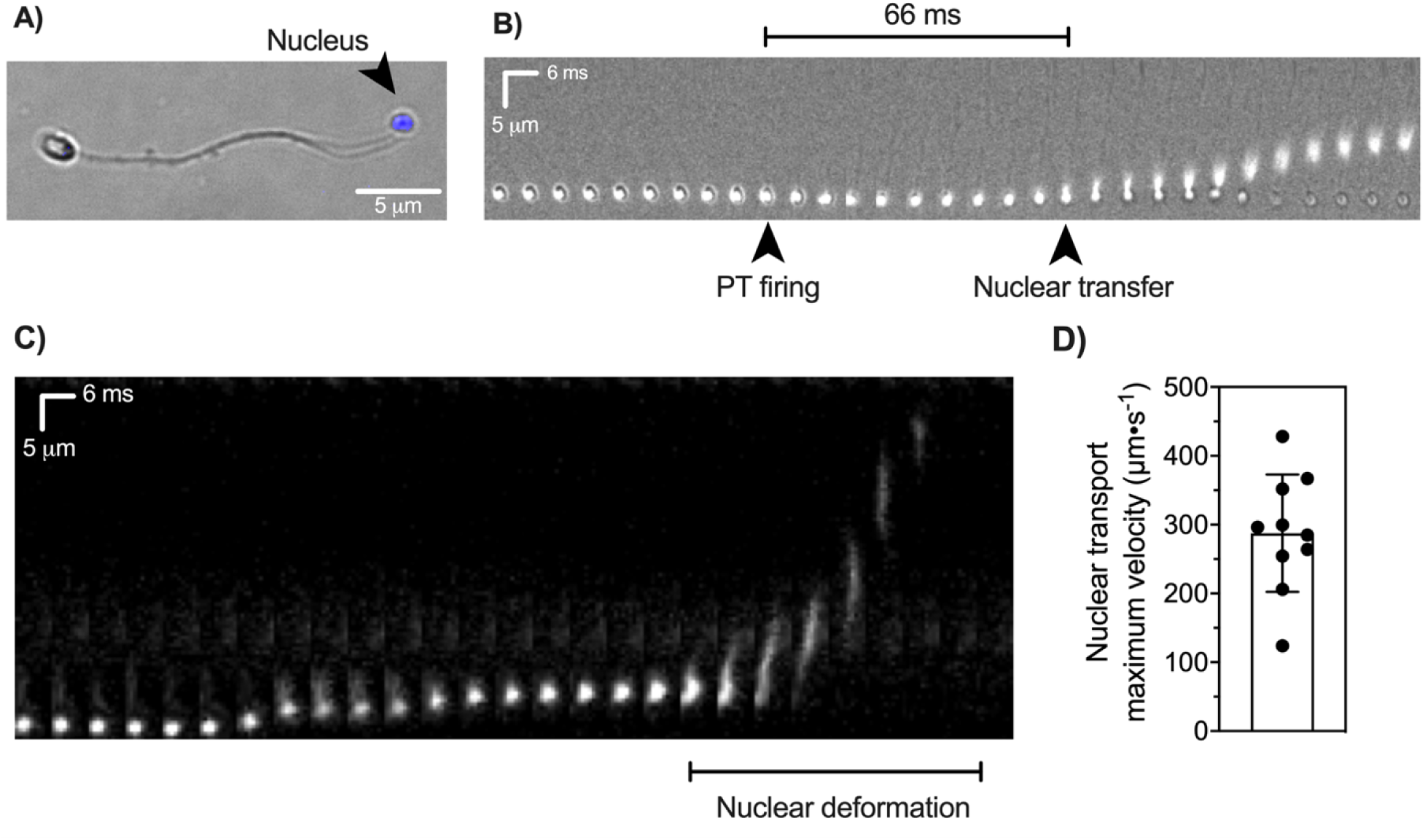
Live-cell imaging of the EHP nuclear transport. (A) A micrograph showing the germinated EHP spores stained with a nuclear staining dye, NucBlue. (B) Time-lapse images of the EHP nuclear translocation inside the spore and during the polar tube extrusion. The time interval for each image is 6 ms. Note that white light was applied to observe the polar tube firing event simultaneously with the nuclear transport. Black arrowheads indicate the frame in which the polar tube firing is initiated and the frame in which the nucleus begins to exit from the spore body. (C) Kymograph of the nuclear translocation through the polar tube. (D) Quantification of the nuclear transport velocity. Error bars represent the standard deviation (n = 10).

## Discussion

Polar tube germination is an important process to initiate microsporidian infection. Conditions used to trigger the spore germination vary greatly among microsporidian species [9]. Here, we characterize *in vitro* germination conditions for the EHP spores. Synthesizing from our data, we describe a mechanistic detail of the EHP polar tube firing process (Fig. 6). First, phthalate, K^+^, and pH 3.0 are required for the EHP spore germination. The optimal temperature for germination ranges between 25-28°C, similar to the shrimp body temperature. These stimuli can penetrate through thick spore wall layers, very quickly (in ∼30 sec). After that, the polar tube extrudes from the apical tip of the spore with the maximum speed of ∼312 μm⋅s^-1^. Once the polar tube is extruded to ∼50% of its length, the infection cargo begins to transport through the tube, similar to what has been previously observed in *A. algerae*. The speed of the cargo transport is comparable to the polar tube firing speed. Massive nuclear deformation is observed in EHP during cargo transport. Eventually (∼117 ms since the polar tube begins to fire), the cargo reaches the end of the polar tube and sporoplasm is clearly visible at the distal end of the polar tube. Furthermore, the ultrastructure of the EHP polar tube shows a membranous layer with an additional repetitive protein unit on the outermost layer of the tube. Further studies are required to understand what proteins constitute the repetitive protein layer of the EHP polar tube and whether the polar tube ultrastructure is conserved among microsporidian species.

**Fig. 6.**
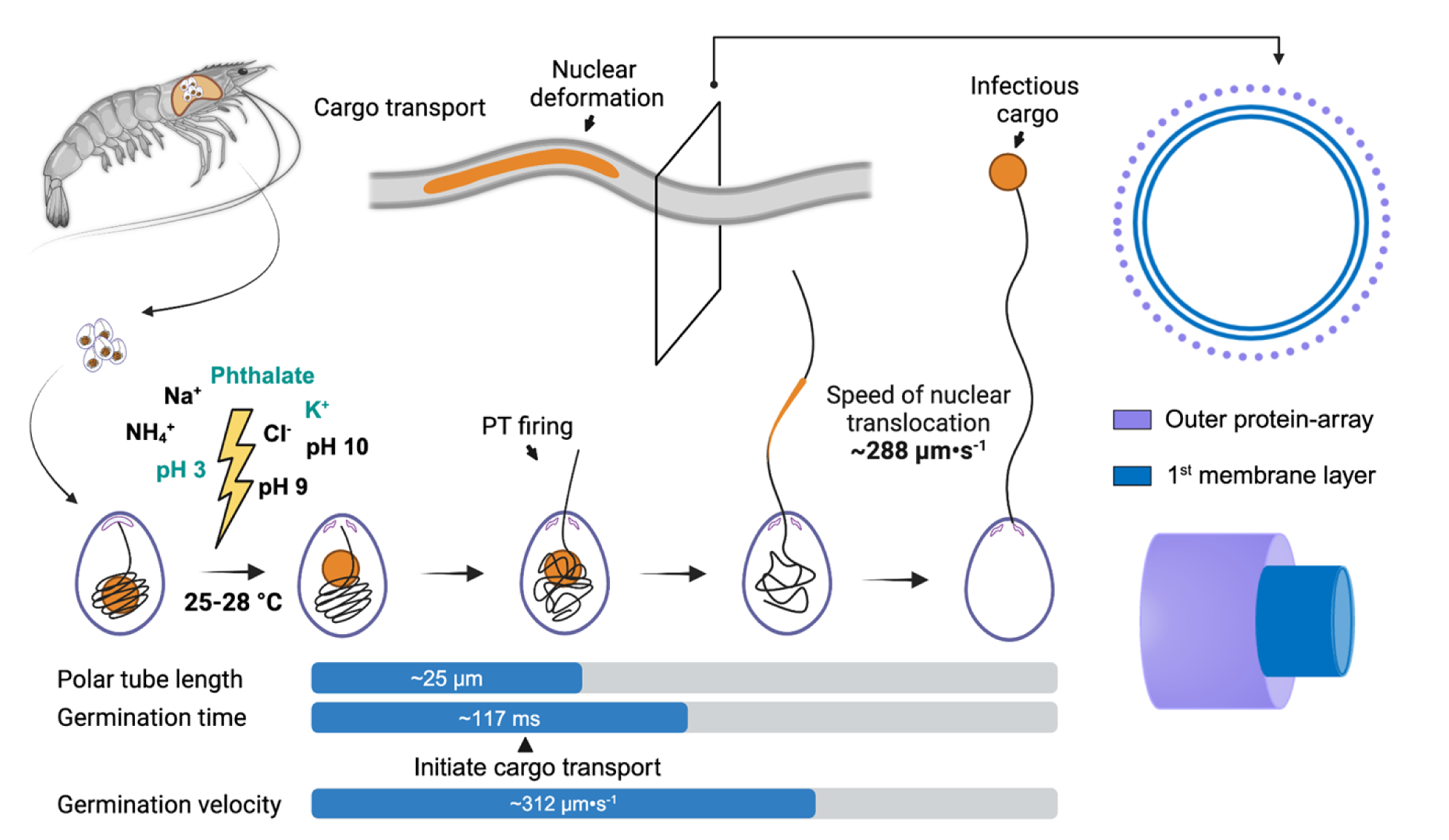
Schematic representation of the EHP polar tube firing process and the ultrastructure of the polar tube. From our data, Phthalate, K^+^, and acidic conditions (pH 3.0) are required for the EHP spore germination, *in vitro*. These stimuli could move across the spore layer in ∼30 sec. The optimal temperature for EHP spore germination ranges between 25-28°C. The polar tube emerges from the apical tip of the spores with a speed of ∼312 μm⋅s^-1^. After the polar tube extends to ∼50% of its total length, the nucleus begins the translocation with comparable speed to the polar tube firing. Ultrastructure of the EHP polar tube by cryoEM revealed two membranes with an extra repetitive protein-array on the surface of the polar tube. These repetitive protein units can possibly be made of PTPs. This figure was created with BioRender.

Physical conditions that trigger spore germination differ greatly depending on microsporidian species. In the genus *Encephalitozoon*, H_2_O_2_ is required for *in vitro* spore germinations [14], whereas in genus *Nosema*, dehydration for a certain period is important for successful spore germination [14]. Little is known about the conditions in the genus *Ecytonucleospora* (*Enterocytozoon*), especially for an economically important species like EHP. Inability to culture spores of this genus *in vitro* has impeded studies involving spore germination. As a result, spores need to be freshly prepared from the infected animals. Our results demonstrate that EHP spores could germinate under both acidic and basic conditions. Ability to germinate under both acidic and basic conditions is less common in microsporidia. So far, two other microsporidian species have been reported to germinate under both conditions, including *Plistophora anguillarum* and *Vavraia culicis*, microsporidian species infecting eels and mosquitoes, respectively [22,23]. For EHP, we showed that both KHP pH 3.0 and the ammonia-ammonium chloride buffer at pH 9.0 or 10.0 could trigger the spore germination. However, the germination rate under an acidic condition was significantly higher. It is plausible that the acidic condition required for the EHP spore germination is due to the prevailing conditions in shrimp digestive tract. There is no direct measurement of pH in each part of the shrimp digestive tract, however several digestive enzymes isolated from penaeid shrimp have their optimal activities in acidic conditions [45]. Alternatively, EHP spore germination might be triggered by a low pH environment in the lysosomal compartment [46]. Since how EHP enters the host cells remains unclear, the spores could be phagocytosed by the cells and later fused with lysosomes, similar to what have been previously observed in *E. cuniculi* spores [47]. In addition to acidic pH, our results demonstrate that EHP spore germination is specific to phthalate molecules. Phthalate is a class of aromatic esters used in several industrial products, such as cosmetics and plastic synthesis [48]. Contamination of phthalate in the shrimp culturing system has been reported in Taiwan [49]. A trace amount of phthalate was detected in two Pacific white shrimp samples (∼0.7 mg/g shrimp) [49]. Further investigations are required to test whether other aromatic compounds could also trigger EHP spore germination. It is important to note that phthalate has been reported to be toxic to the environment and human health by causing respiratory and liver cancers, even at 150 mg/kg/day [50,51]. The use of phthalate as an anti-EHP agent needs to be tested in the laboratory and the farm settings to mitigate the associated risk to human health and the environment.

Polar tube is a specialized invasion organelle conserved across all microsporidian species [52]. The ultrastructures of the microsporidian polar tube have been previously described in several microsporidian species. Cryo-EM of the *N. bombycis* polar tube showed repetitive bumps on the surface of the extruded tube with a distance of ∼2 nm between each unit, resembling comb teeth [35]. It is possible that these bumps might be PTPs. In contrast to *A. algerae*, the outermost layer of the extruded polar tube is covered by fine fibrillar materials, possibly due to glycosylation [53]. The differences in the polar tube structure might be due to different microsporidian species or stages of the tube whether they contain cargo or not. For EHP, we observe repetitive protein units on the outermost layer of the extruded polar tube. The distance between each protein unit is ∼52-66 Å as quantified from 2D class averaging. Similar spacing was observed in the outer filament (OF)-layer of the *E. intestinalis* polar tube (∼58 Å) [39]. It is suggested that the repeating units along the filaments may be composed of one or more PTPs [39]. It is interesting that these ∼58 Å repetitive units are found in the interior part of the tube prior to firing in *E. intestinalis*, while the repetitive units are on the outermost surface of the extruded polar tube in EHP. With an assumption that if the proteins made of these repetitive units are conserved between *E. intestinalis* and EHP, these results would support the eversion model that the tube is turning inside out during the germination process. However, direct comparison of the distance between repetitive units on the same microsporidian species, pre- and post-firing is needed to further support this eversion model.

## Materials and methods

### Propagation of EHP

Specific pathogen-free (SPF) *Litopenaeus vannamei* juveniles were provided by the Marine Shrimp Broodstock Research Center II (MSBRC-2), Charoen Pokphand Foods PCL (Phetchaburi Province, Thailand). EHP-infected shrimps were obtained from commercial ponds in Chanthaburi province, Eastern Thailand for all germination analyses and from the Aquaculture Pathology Laboratory, Tucson, University of Arizona, USA for cryoEM analysis. To propagate the parasites in shrimp, SPF shrimps were reared with EHP-positive shrimps by co-habitation method for fourteen days, as previously described [54]. Mature EHP spores were purified by a discontinuous Percoll gradient [55]. In brief, hepatopancreas from 10-15 shrimps were homogenized by a glass pressure homogenizer. The cell lysates were filtered with a 40-μm cell strainer (Jet Biofil, China) and centrifuged with 50% Percoll (1:1 ratio). The mature spores were further isolated using 0, 25, 50, 75, and 100% of Percoll solution [55]. EHP mature spores were washed and stored in 1X PBS for further use.

### Identification of optimal conditions for EHP spore germination

Previous studies have shown important factors essential for spore germination, such as a shift of pH [17], alkali metal ions [18], and high salt concentrations [18]. To identify the optimal condition for EHP spore germination, two μl of EHP mature spores (10^8^ spores/ml) were incubated with various germination buffers at different pH, ranging from pH 2.0-13.0, The buffers include KCl (pH 2.0, 9.5, 12.0, and 13.0), K_3_C_6_H_5_O_7_ (pH 3.0, 4.0, 5.0, 6.0), KHP (pH 2.0, 3.0, and 4.0), KH_2_PO_4_ (pH 6.0, 7.0, and 8.0), Borax (pH 9.5), Glycine (pH 9.0 and 10.0), NH_3_/NH_4_Cl buffer (pH 9.0 and 10.0), NaHCO_3_ (pH 10.0 and 11.0), Na_2_HPO_4_ (pH 11.0 and 12.0) and NaOH (pH 13.0). To further examine the optimal concentration of the germination buffers, spores were incubated with 25, 50, 75, and 100 mM of the germination buffers. We further investigated the effect of KCl concentrations by incubating the spores at 100, 200, 300, 400, and 500 mM concentrations. Temperature and time are important factors that influence spore germination [26,56]. The mature spores were incubated at 10, 25, 28, 37 and 40°C. Furthermore, we varied the incubation time to study its impact on the germination. The incubation times used in this study were 0.5, 1, 5, 10, 20, and 30 minutes. For spore examination under each condition, spores were incubated for 30 minutes at room temperature, except in the test assessing the effect of time, where the durations were varied. For the reactions of temperature and time experiments, the spores were fixed with 4% formaldehyde for 30 min prior to accessing the germination rate. The spores were later placed onto a poly-L-lysine-coated slide (Huida, China) and sealed with a coverslip. Then, they were observed under an inverted microscope (Zeiss AxioObserver) using a 60X oil objective lens. The images were captured in consecutive fields, and the polar tube extrusions were counted for at least 100 spores per germination condition. Each experiment was performed in three biological replicates.

### Live-cell imaging of the EHP polar tube firing process

The polar tube firing happens on an extremely fast timescale (less than 2 sec) [26]. Due to its fast nature, high-speed live-cell imaging is used to better understand the kinetics of the polar tube firing. To trigger the polar tube firing process in EHP, two μl of purified EHP spores (10^8^ spores/ml) were mixed with 2% (w/v) phloxine B solution or with 100 mM potassium hydrogen phthalate (KHP) pH 3.0. Then, the reaction was placed on a poly-L-lysine-coated slide and sealed with a #1.5 coverslip. The slide is quickly placed on a Zeiss AxioObserver inverted microscope with a 60X oil immersion objective lens. The experiment was performed at room temperature, which is suitable for EHP spore germination. The exposure time for recording the video was adjusted to maximize the frame rates. We could drive the microscope to ∼170 frames per second. The videos were collected from at least 20 different EHP spores to see the variation of the polar tube firing kinetics. After obtaining the polar tube firing videos, they were processed using ImageJ [57]. Kymographs were generated using a straighten function. The polar tube length at different time intervals was traced and measured using a segmented line function. The polar tube firing velocities and accelerations were calculated by Δy/Δx where Δy is change in polar tube length or change in velocities, and Δx is change in time intervals. Graphs were generated using GraphPad software.

### Live-cell imaging of the nuclear transport

To observe the cargo transport through the polar tube, 100 μl of purified EHP spores (10^8^ spores/ml) were mixed with 1 drop of NucBlue (Invitrogen, catalog no. R37605) at room temperature for 10 min. Then, 2 μl of the pre-stained spores were mixed with 10 μl of the potassium phthalate buffer, pH 3.0. The reaction was immediately placed on a poly-L-lysine-coated glass slide and covered with a #1.5 18 x 18 mm coverslip. The imaging was performed using a Zeiss AxioObserver inverted microscope with 60X oil immersion objective lens. The camera provided a wide field of view at 273 frames per second with 4 ms exposure time and 30% laser intensity. 2X2 binning was also applied.

### Cryo-EM sample preparation and data collection

The crude EHP spores used for CryoEM were isolated EHP-infected *P. vannamei* at Aquaculture Pathology Laboratory, The University of Arizona. Spores were kept in a thermos to control the temperature at ∼RT during the shipment. Upon arrival at the Bhabha/Ekiert laboratory at Johns Hopkins University, the EHP mature spores were isolated as described in the previous section. We tested the germination rate of the spores upon arrival by incubating the spores with 2% phloxine B and 100 mM KHP pH 3.0 for 5 min at room temperature. The germination rate was ∼20% in both KHP and phloxine B induced conditions. To visualize the EHP polar tube architecture using CryoEM, the EHP polar tube firing was triggered using two different germination buffers, including 2% phloxine B and 100 mM KHP. In this experiment, 15 μl of the mature EHP spores were incubated with 75 μl of 100 mM KHP buffer, pH 3.0 or with 2% (w/v) phloxine B for 5 min at room temperature. Then, the spores were placed on a lacey carbon film with 300 mesh (Catalog number: LC300-Cu-150) (Electron Microscopy Sciences, USA), followed by a C-SPAM vitrification system [58] with a blotting time of 6 sec prior to plunge freezing with liquid ethane. Note that in the KHP induced condition, the purified EHP spores were kept at room temperature for 9 days prior to blotting. The germination rate on Day 9 was still ∼20%. Moreover, the germinated spores were washed with 1X PBS pH 7.4, followed by centrifugation at 200 xg for 5 min prior to being placed on an EM grid. The grids were imaged on a 300-kV Titan Krios G3i transmission electron microscope (Thermo Fisher Scientific, USA) at the Beckman Center for CryoEM at Johns Hopkins, using a Falcon 4i direct electron detector (Thermo Fisher Scientific, USA) with a magnification of 81,000X, corresponding to a calibrated pixel size of 1.23 Å/pixel with total dose of 29.45 e/Å^2^ and the defocus at −1.8 to −2.3 μm.

### Cryo-EM data processing

The cryoEM data processing workflow is summarized in S2 Fig. In brief, movies were imported into cryoSPARC v4.6 [59]. Movies were corrected using the patch motion correction tool within cryosparc, and CTF was determined via the patch CTF estimation job. Initial particle picking was done with the filament tracer tool, leading to 13,868 particles were picked from 112 micrographs in the KHP condition, and 11,683 particles were picked from 104 micrographs under the phloxine-B condition. Micrographs were curated to remove those where the tube appears to be broken and the presence of ice crystals is evident, and particle picking adjusted using the thresholds of the template picker tool. This process resulted in 2,061 particles in phloxine B and 3,701 particles in the KHP conditions. Those particles were extracted using a box size of 350 pixels, and used for a first round of 2D classification. A second round of 2D classification was performed by selecting only subclasses in which the particles were localized on the outer layer of the tube. A final round of 2D classification was conducted to categorize the particles with similar patterns together. These processes allowed us to obtain two groups for the phloxine B sample and one for the KHP sample.

These three groups show a lipid bi-layer with regular densities likely representing organized protein structure protruding outside the polar tube. Those protein structures seem to show different organizations in the 3 groups and to characterize them, the 2D classes images were imported into ImageJ [57]. The grey level profiles along and across the protein patterns were drawn to measure the distance between the protein densities and the distance to the membrane. More precisely, the distance between protein densities and the width of the membrane bi-layer were measured from peak-to-peak profile densities along the tube axis (S4A Fig and S4B Fig).

Finally, the particles belonging to each class were projected back to their micrograph of origin, and polar tube sections un-ambiguously enriched in particles from a given group were associated with that group. The polar tube diameters in regions associated with particles from each group were then measured from one side of the repetitive protein unit to another.

### Statistical analyses

All statistical analyses were conducted using GraphPad Prism 9 software. For the physical condition analysis, one-way ANOVA was used for Fig. 1B, 1C, and 1D, while a two-tailed paired Student’s t-test was used in Fig. 1E and 1F. For the quantification of kinetic parameters, one-way ANOVA was used to analyze the differences between conditions. For the protein organization of PT, we used a two-tailed unpaired Student’s t-test to compare differences between two groups.

## Acknowledgments

CryoEM data were collected at the Beckman Center for CryoEM at Johns Hopkins. We thank Kai Cai, Dazhong Ding, and Duncan Sousa for their assistance with project planning and tomography data collection. This processing of the cryoEM data was carried out at the Advanced Research Computing at Hopkins (ARCH) core facility (rockfish.jhu.edu), which is supported by the National Science Foundation (NSF) grant number OAC1920103.

## Supplementary information

**S1 Fig.**
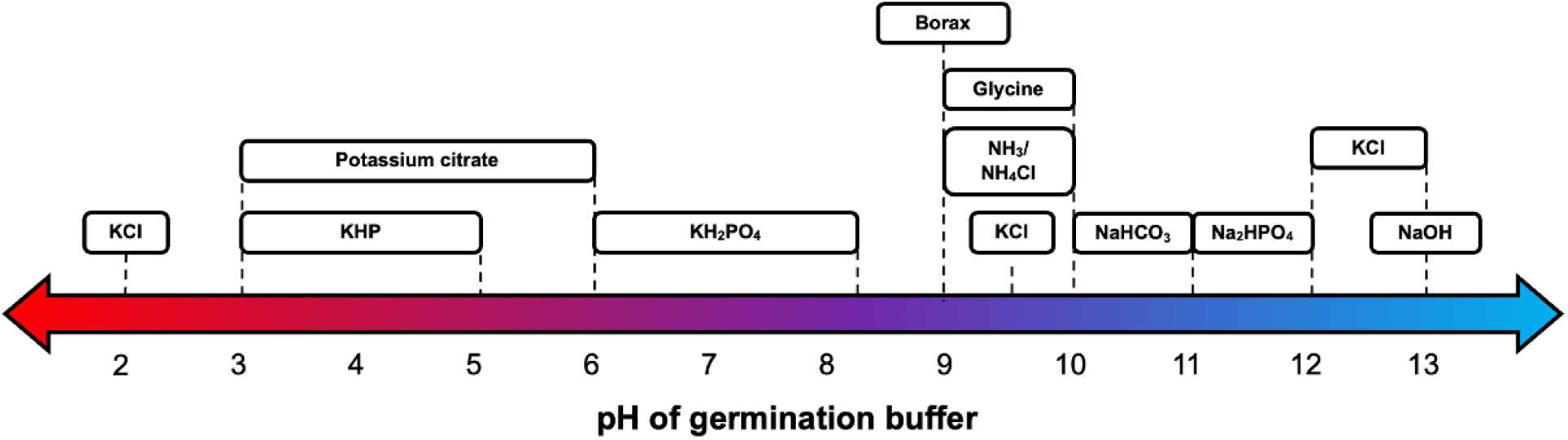
Germination buffers used to test EHP spore germination, *in vitro*. Difference germination buffers and pHs were used to trigger the EHP spore germination, including KCl (pH 2.0, 9.5, 12.0, and 13.0), K_3_C_6_H_5_O_7_ (pH 3.0, 4.0, 5.0, 6.0), KHP (pH 2.0, 3.0, and 4.0), KH_2_PO_4_ (pH 6.0, 7.0, and 8.0), Borax (pH 9.5), Glycine (pH 9.0 and 10.0), NH_3_/NH_4_Cl (pH 9.0 and 10.0), NaHCO_3_ (pH 10.0 and 11.0), Na_2_HPO_4_ (pH 11.0 and 12.0) and NaOH (pH 13.0).

**S2 Fig.**
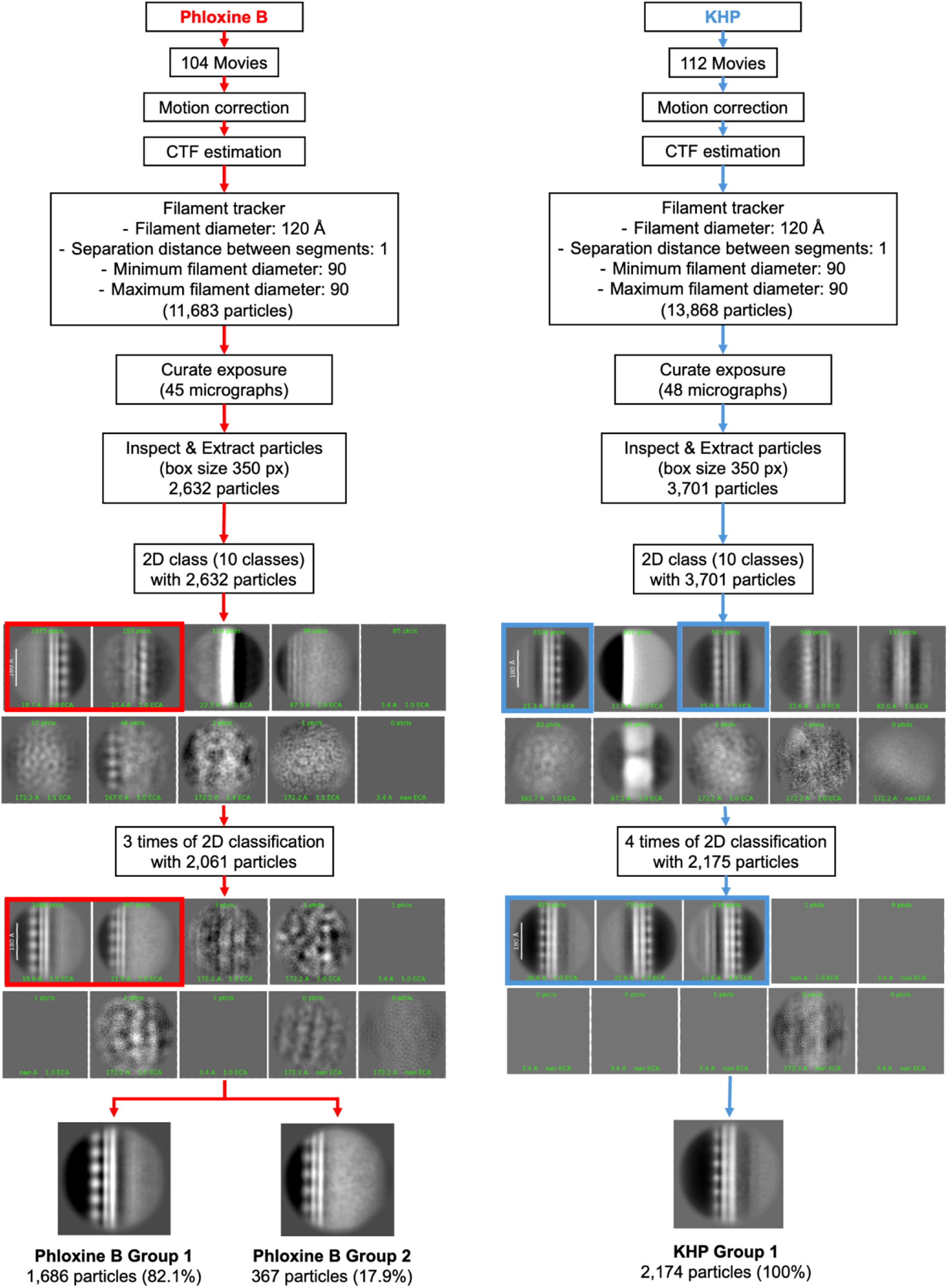
CryoEM data processing workflow. A cryo-EM data processing workflow to visualize the EHP polar tube architecture when the spores were induced with phloxine B (left) and KHP (right). Movies were imported into CryoSPARC. They were aligned using patch motion correction and patch CTF, followed by tracking the filament at the edge of the polar tubes. The particles were picked and extracted to 350 pixels. Then, the 2D classification was performed.

**S3 Fig.**
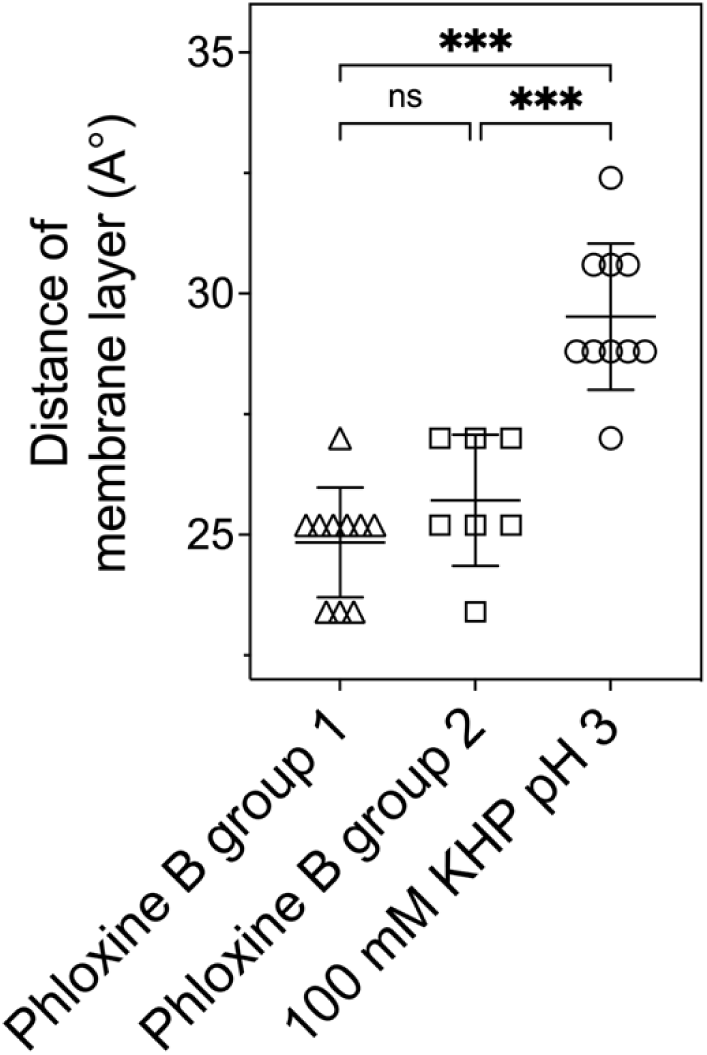
The thickness of the membranous layer from three subclasses of 2D class averaging.

**S4 Fig.**
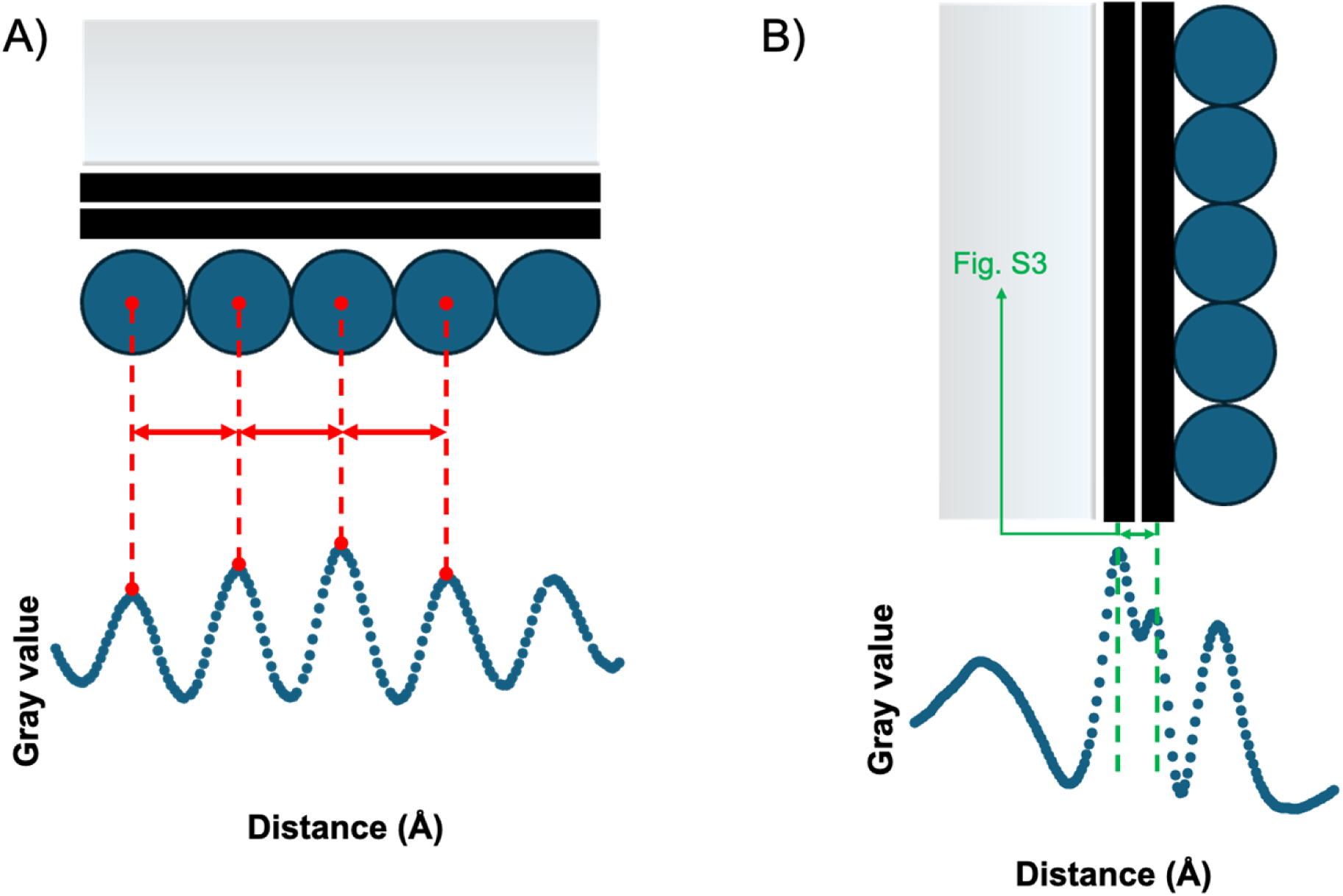
Quantification of protein subunit from 2D class averaging. (A) Measurement of the spacing between each repetitive protein and (B) The width of the membrane bilayer.

**S1 Table.**
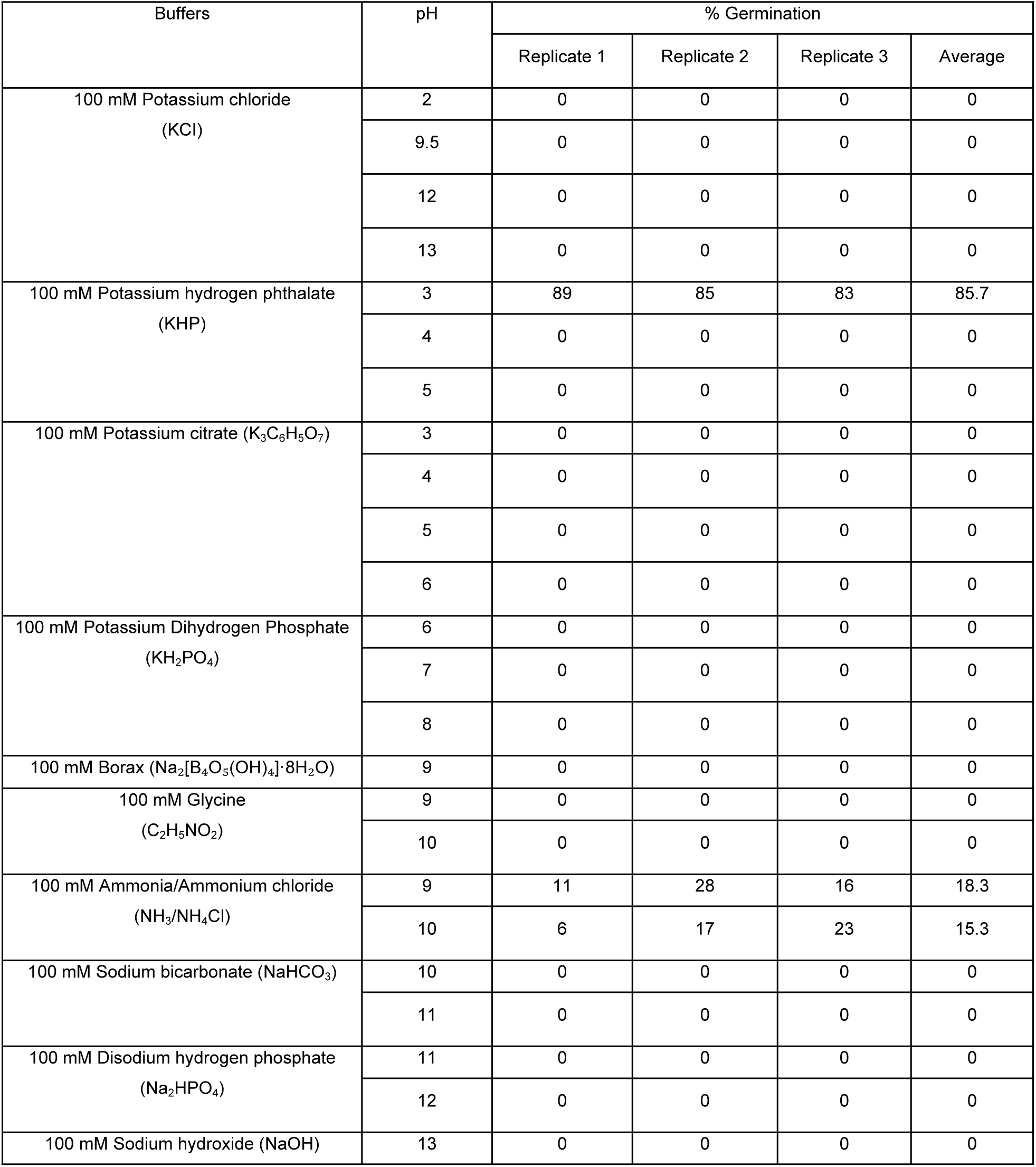
Quantification of the EHP germination rate in different germination buffers.

## Supplementary videos

**S1 Video.** Live-cell imaging of the EHP spore germination induced by 100 mM KHP pH 3 at 25°C

**S2 Video.** Live-cell imaging of the EHP spore germination induced by 100 mM KHP pH 3 at 37°C

**S3 Video.** Live-cell imaging of the EHP spore germination induced by 2% phloxine B

**S4 Video.** Visualization of the polar tube firing together with a nuclear translocation

**S5 Video.** Live-cell imaging of the EHP nuclear transport

